# Coiled-coil registry shifts in the F684I mutant of Bicaudal result in cargo-independent activation of dynein motility

**DOI:** 10.1101/776187

**Authors:** Heying Cui, Kathleen M. Trybus, M. Yusuf Ali, Puja Goyal, Kaiqi Zhang, Jia Ying Loh, Sozanne R. Solmaz

## Abstract

The dynein adaptor *Drosophila* Bicaudal D (BicD) is auto-inhibited and activates dynein motility only after cargo is bound, but the underlying mechanism is elusive. In contrast, we show that the full-length BicD/F684I mutant activates dynein processivity even in the absence of cargo. Our X-ray structure of the C-terminal domain of the BicD/F684I mutant reveals a coiled-coil registry shift; in the N-terminal region, the two helices of the homodimer are aligned, whereas they are vertically shifted in the wild-type. One chain is partially disordered and this structural flexibility is confirmed by computations, which reveal that the mutant transitions back and forth between the two registries. We propose that a coiled-coil registry shift upon cargo binding activates BicD for dynein recruitment. Moreover, the human homolog BicD2/F743I exhibits diminished binding of cargo adaptor Nup358, implying that a coiled-coil registry shift may be a mechanism to modulate cargo selection for BicD2–dependent transport pathways.

## INTRODUCTION

Cytoplasmic dynein, the predominant minus-end directed microtubule motor, facilitates a vast number of cellular transport events (Cianfrocco *et al.*, 2015). Dynein adaptor proteins, such as *Drosophila* Bicaudal (*Dm* BicD) (Hoogenraad *et al.*, 2001) recognize cargo for dynein-dependent transport. Cargo-bound adaptors are required to activate metazoan dynein for processive transport and are therefore an essential part of the dynein transport machinery (Chowdhury *et al.*, 2015; McKenney *et al.*, 2014; Schlager, Hoang, *et al.*, 2014; Schlager, Serra-Marques, *et al.*, 2014; Splinter *et al.*, 2012; Urnavicius *et al.*, 2015). In the absence of cargo, BicD forms an auto-inhibited looped conformation, in which the C-terminal cargo binding region (CTD) binds to the N-terminal dynein/dynactin-binding site (coiled-coil domain 1, CC1), sterically preventing motor binding (Chowdhury *et al.*, 2015; Liu *et al.*, 2013; McClintock *et al.*, 2018; McKenney *et al.*, 2014; Schlager, Hoang, *et al.*, 2014; Schlager, Serra-Marques, *et al.*, 2014; Sladewski *et al.*, 2018; Splinter *et al.*, 2012; Terawaki *et al.*, 2015; Urnavicius *et al.*, 2015). The CTD is required for auto-inhibition, since a truncated BicD-CC1 construct without it activates dynein for processive transport in the absence of cargo (McKenney *et al.*, 2014; Schlager, Hoang, *et al.*, 2014). Auto-inhibition of full-length BicD is released upon cargo binding (Chowdhury *et al.*, 2015; Liu *et al.*, 2013; McClintock *et al.*, 2018; McKenney *et al.*, 2014; Schlager, Hoang, *et al.*, 2014; Schlager, Serra-Marques, *et al.*, 2014; Sladewski *et al.*, 2018; Splinter *et al.*, 2012; Terawaki *et al.*, 2015; Urnavicius *et al.*, 2015); however, the underlying molecular mechanism is elusive.

Notably, auto-inhibition is compromised in the classical *Dm* BicD Bicaudal mutant F684I, which recruits larger amounts of dynein from cell extracts compared to wild-type *Dm* BicD (Liu *et al.*, 2013). The mutation causes dynein and Egalitarian-dependent mRNA transport defects and subsequent anterior accumulation of the *Oskar* mRNA pool (Mach & Lehmann, 1997; Mohler & Wieschaus, 1986; Navarro *et al.*, 2004; Zimyanin *et al.*, 2008), which result in a classical developmental phenotype that includes double-abdomen flies (Bull, 1966; Liu *et al.*, 2013; Mohler & Wieschaus, 1986; Wharton & Struhl, 1989). Therefore, the Bicaudal mutant could potentially serve as a tool to investigate the molecular mechanism of BicD-auto-inhibition.

*Dm* BicD facilitates the transport of mRNA and Golgi-derived vesicles and is recruited to these cargoes via protein cofactors that are termed cargo adaptors. The most well characterized cargo adaptors for *Dm* BicD are Egalitarian (Dienstbier *et al.*, 2009) and Rab6^GTP^, which facilitate transport of mRNAs and Golgi-derived vesicles, respectively (Matanis *et al.*, 2002). The predominant cargo adaptors for the human homologs Bicaudal D2 (*Hs* BicD2) and Bicaudal D1 (*Hs* BicD1) are Rab6^GTP^ (Matanis *et al.*, 2002), which engages in the transport of Golgi-derived and secretory vesicles, and nuclear pore complex protein Nup358, which engages in transport of the cell nucleus (Splinter *et al.*, 2010). BicD2-dependent transport pathways are important for faithful chromosome segregation, neurotransmission at synapses, and essential for brain development (Baffet *et al.*, 2015; Hu *et al.*, 2013; Splinter *et al.*, 2010). Mutations of BicD2 cause the neuromuscular disease spinal muscular atrophy (Martinez-Carrera & Wirth, 2015; Peeters *et al.*, 2013; Synofzik *et al.*, 2014). Cargo selection for BicD2-dependent transport is regulated by competition of cargo-adaptors (Noell *et al.*, 2018), as well as the G2 phase specific kinase Cyclin-dependent kinase 1 (Cdk1) (Baffet *et al.*, 2015), however, additional regulatory mechanisms remain to be identified in order to explain how BicD2 switches from selecting Rab6-positive vesicles for transport in G1/S phase to recruiting dynein to the cell nucleus via Nup358 in G2 phase.

Cargo adaptors bind to the C-terminal domain (CTD) of *Dm* BicD (Hoogenraad *et al.*, 2001), and the structure of the *Dm* BicD-CTD (Liu *et al.*, 2013) as well as its homologs *Hs* BicD2 (Noell *et al.*, 2019) and mouse (*Ms*) BicD1 (Terawaki *et al.*, 2015) were determined, which all form homodimeric coiled-coils. Interestingly, *Hs* BicD2 and *Ms* BicD1 form a conformation with homotypic coiled-coil registry, in which the helices are aligned at equal height and the same residues from both chains engage in layers of knobs-into-holes interactions (Noell *et al.*, 2019; Terawaki *et al.*, 2015). In contrast, *Dm* BicD has an asymmetric coiled-coil registry. In the N-terminal half of the CTD, the helices are vertically shifted by ∼1 helical turn respective to each other (heterotypic coiled-coil registry), whereas in the C-terminal half, the chains are aligned in a homotypic coiled-coil registry (Liu *et al.*, 2013). Coiled-coil registry shifts have so far only been reported for a few proteins, including dynein (Carter *et al.*, 2008; Choi *et al.*, 2011; Croasdale *et al.*, 2011; Gibbons *et al.*, 2005; Kon *et al.*, 2009; Macheboeuf *et al.*, 2011; Noell *et al.*, 2019; Snoberger *et al.*, 2018; Stathopulos *et al.*, 2013; Xi *et al.*, 2012), but may potentially be an inherent property of many coiled-coil structures with important physiological functions. In the case of BicD2, a coiled-coil registry shift may relieve auto-inhibition.

We recently used molecular dynamics (MD) simulations to probe structural dynamics in the BicD2-CTD coiled-coil (Noell *et al.*, 2019). These simulations support the idea that BicD2 can adopt both a homotypic coiled-coil registry, and an asymmetric registry, as both states are similarly stable in simulations and defined by distinct conformations of F743 and F750, which stabilize either a homotypic or asymmetric coiled-coil registry (Noell *et al.*, 2019). Notably, mutation of F743 to Ile (F684I in *Dm*) increases dynein recruitment in the *Drosophila* homolog compared to the wild type (Liu *et al.*, 2013). In our MD simulations of the F743I mutant of *Hs* BicD2-CTD, a spontaneous coiled-coil registry shift from asymmetric to fully heterotypic coiled-coil registry was observed (Noell *et al.*, 2019). We thus hypothesized that a coiled-coil registry shift upon cargo binding could relieve BicD-auto-inhibition and activate it for dynein recruitment, as has also been proposed earlier (Liu *et al.*, 2013; Noell *et al.*, 2019; Terawaki *et al.*, 2015). In addition, in MD simulations of the R747C human disease mutant of *Hs* BicD2-CTD which causes spinal muscular atrophy, a spontaneous transient coiled-coil registry shift was observed, which may be an underlying cause of the disease (Noell *et al.*, 2019).

Here we show that the Bicaudal mutation F684I abolishes auto-inhibition and allows cargo-independent activation of dynein motility. To investigate the structural basis for this activation, we determined the X-ray structure of the C-terminal cargo-binding domain (CTD) of *Dm* BicD-CTD/F684I, which has a homotypic coiled-coil registry, in contrast to the wild-type. Furthermore, in the structure of the mutant, the region N-terminal of F684I is disordered for one chain. This structural flexibility is confirmed by MD simulations, in which the mutant transitions back and forth between homotypic and asymmetric registries on a time scale of tens of ns. Free energy calculations indicate conformations with homotypic and asymmetric registries to have similar stability. Our data suggest that the mutation promotes a registry shift and renders the coiled-coil flexible, likely resulting in the formation of multiple conformations.

## RESULTS

### Full-length Dm BicD with the F684I mutation recruits dynein in the absence of cargo

A single molecule TIRF (Total Internal Reflection Fluorescence) microscopy processivity assay was used to assess the functional properties of reconstituted dynein-dynactin-BicD (DDB) complexes. Complexes were reconstituted with either full-length *Dm* BicD (BicD^WT^), full-length *Dm* BicD/F684I (BicD^F684I^), or the truncated N-terminal fragment of *Dm* BicD (BicD^CC1^). The molar ratio of dynein-dynactin:BicD was 1:1:2 to ensure recruitment of only 1 dynein to the ternary complex (Sladewski *et al.*, 2018). Full-length WT *Dm* BicD is auto-inhibited and does not recruit dynein-dynactin, while the truncated N-terminal fragment fully activates dynein-dynactin for processive transport (McClintock *et al.*, 2018; McKenney *et al.*, 2014; Schlager, Hoang, *et al.*, 2014; Sladewski *et al.*, 2018). Consistent with previous results (Sladewski *et al.*, 2018), we did not observe processive directional movement for DDB^WT^ on microtubule tracks, although some one-dimensional diffusive events of the Qdot-labeled dynein were observed. Because DDB^WT^ showed no directional motion (Movie S1), speed and run length were not measured. In contrast, DDB^CC1^ showed robust processivity, which was indistinguishable from that observed with DDB^F684I^ (Fig. 1A,B, Movie S2 and S3). The speeds of both DDB^CC1^ and DDB^F684I^ were fitted with a single Gaussian (Fig. 1C). The DDB^F684I^ complex moved at a speed of 0.43 ± 0.17 µm/s (n=68), which was not significantly different (*p*=0.44) from what was observed for DDB^CC1^ (0.41 ± 0.21 µm/s, n=58). Processive run-lengths were fitted with a standard exponential decay equation (y= Ae^-bx^), where A is the amplitude and 1/b is run length. The run length of DDB^F684I^ (3.3 ± 0.18 µm, n=68) was not significantly different (*p*=0.9) from that of DDB^CC1^ (2.8 ± 0.13 µm, n=58) (Fig. 1D).

**Figure 1.**
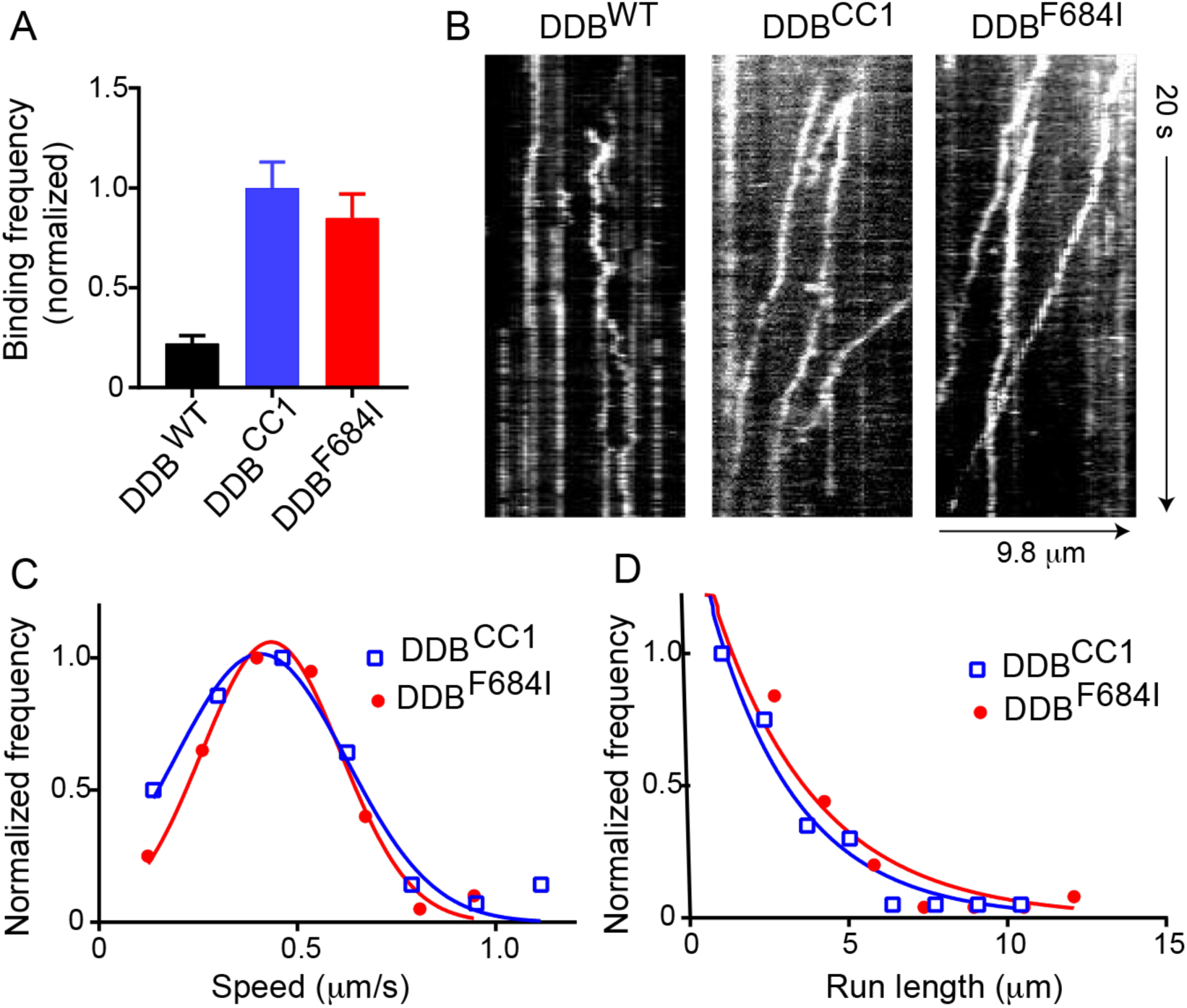
Full-length *Dm* BicD^F684I^ results in cargo-independent activation of dynein motility. (A) Normalized run frequency of dynein-dynactin-BicD (DDB) with different *Dm* BicD constructs: full-length BicD^WT^ (black), truncated BicD^CC1^ (blue), and full-length mutant BicD^F684I^ (red). The binding frequency of DDB^CC1^ is normalized to one. Frequencies of DDB^CC1^ and DDB^F684I^ are 4.5-fold and 3.8-fold higher, respectively, than DDB^WT^. Motion of DDB^WT^ is mainly diffusive. (B) Kymographs of the three DDB complexes. DDB^WT^ is either static or diffusive, while the other two complexes show processive motion (sloped lines). (C) The speed of DDB^CC1^ (blue) and DDB^F684I^ (red) are 0.41 ± 0.21 µm/s, n=58) and 0.43 ± 0.17 µm/s (n=68) (mean ± SD), respectively, which were not significantly different (*p*=0.44). (D) Run length of DDB^CC1^ (2.8 ± 0.13 µm, n=58, blue) and DDB^F684I^ (3.3 ± 0.18 µm, n=68, red) (mean ± SD) were not significantly different (*p*=0.9). Data from 3 independent experiments were pooled.

Based on run frequency, speed and processive run length, we conclude that the full-length mutant *Dm* BicD^F684I^ was not auto-inhibited and was fully capable of binding and activating dynein-dynactin for processive transport in the absence of cargo.

### Crystal structure of the Dm BicD-CTD/F684I mutant provides mechanistic insights into cargo-independent activation

To gain structural insights into the molecular mechanism of activation of BicD, we determined the structure of the C-terminal domain of the *Drosophila melanogaster* BicD/F684I mutant (*Dm* BicD-CTD/F684I, residues 656-745). Crystals were obtained in the space group *P*3_1_ 2 1. The structure was determined by molecular replacement in the PHENIX suite (Adams *et al.*, 2010), using coordinates from the wild-type structure (Liu *et al.*, 2013) that were truncated N-terminal of residue 692 as the search model. The structure was refined to 2.35Å resolution, with an R_free_ of 25.99% and an R_work_ of 25.06% (Table 1). In the structure, *Dm* BicD-CTD/F684I forms a homodimeric coiled-coil. However, in contrast to the structure of the wild type, in the structure of the Bicaudal mutant, a ∼20 residue N-terminal region upstream of the mutated residue I684 is not resolved in the electron density map for one chain of the dimer (Fig. 2). However, the same region is well defined in the electron density map for the second chain. Consequently, the model contains residues 666-740 and 684-741 for the two chains, respectively.

**Table 1.** Crystallographic statistics.

|  |  |
| --- | --- |
| **Data collection statistics** |  |
| Space group | $P 3_1 2 1$ |
| Unit cell parameters a, b, c (Å) | 60.0, 60.0, 142.6 |
| Unit cell parameters $\alpha$, $\beta$, $\gamma$ | 90°, 90°, 120° |
| Wavelength (Å) | 0.9791 |
| Resolution range (Å) | 19.47-2.35 (2.434-2.35) |
| Total Reflections | 125766 (12111) |
| Unique Reflections | 12955 (1263) |
| Redundancy | 9.7 (9.6) |
| Completeness (%) | 99.39 (99.92) |
| $I/\sigma(I)$ | 27.98 (1.26) |
| Wilson B-factor (Å<sup>2</sup>) | 72.37 |
| $R_{\text{merge}}$ | 0.0443 (1.621) |
| $R_{\text{meas}}$ | 0.04684 (1.713) |
| $R_{\text{pim}}$ | 0.01498 (0.549) |
| CC1/2 | 1 (0.573) |
| CC* | 1 (0.854) |
| **Refinement statistics** |  |
| $R_{\text{free}}$ (%) | 25.99 (38.38) |
| $R_{\text{work}}$ (%) | 25.06 (36.04) |
| Reflections (work/test sets) | 12948/640 |
| R.m.s.d. bonds (Å)/angles | 0.005/0.77 |
| Average B-factor (Å<sup>2</sup>) | 96.46 |
| **MolProbity validation** |  |
| Ramachandran plot favored | 100% |
| Ramachandran plot allowed | 0% |
| Ramachandran plot outliers | 0% |
| Rotamer outliers | 0.85% |
| Clash score | 1.83 |

The statistics for the high-resolution shell are shown in parenthesis.

**Figure 2.**
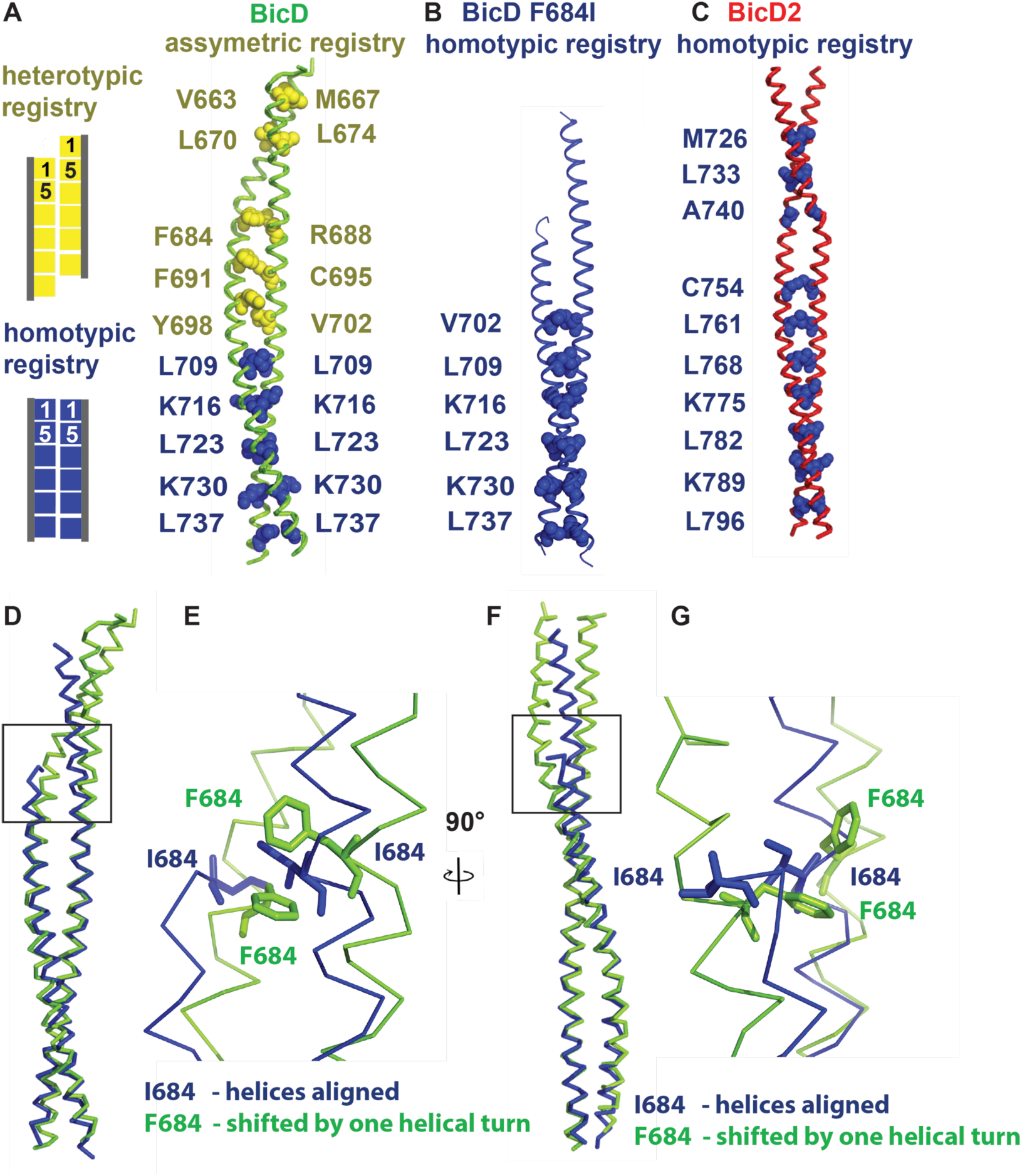
*Dm* BicD-CTD/F684I assumes a conformation with homotypic coiled-coil registry. (A) The structure of *Dm* BicD-CTD wild type (PDB ID 4BL6) (Liu *et al.*, 2013) which has an asymmetric coiled-coil registry is shown in cartoon representation next to a schematic illustrating coiled-coil registries (left panel). Knob residues in the “a” position of the heptad repeat are shown in spheres representation (heterotypic registry yellow, homotypic registry dark blue). (B) Structure of *Dm* BicD-CTD/F684I, which has a homotypic registry. (C) Structure of *Hs* BicD2-CTD (PDB ID 6OFP) (Noell *et al.*, 2019), which has a homotypic registry. (D-G) Least squares superimposed structures of the *Dm* BicD-CTD wild type (green) and the F684I mutant (dark blue) are shown as C-α traces, and are rotated by 90° in (D, F). (E, G) Close-up of the boxed area in (D, F). Residues F684 and I684 are shown in stick representation. Note that in the structure of the Bicaudal mutant, the I684 residues from both chains of the dimer are aligned at the same height, consistent with a homotypic registry, while in the wild-type structure, the F684 residues from both monomers are vertically shifted by one helical turn respective to each other, consistent with a heterotypic registry.

Coiled-coils such as *Dm* BicD-CTD are characterized by heptad repeats ‘abcdefg’ in the sequence, where residues at ‘a’ and ‘d’ positions are predominantly hydrophobic. These residues form characteristic “knobs-into-holes interactions”, where a knob from one chain (either an ‘a’ or ‘d’ position residue) fits into a hole formed of four residues on the opposite chain (Crick, 1953; O’Shea *et al.*, 1991). Notably, structures of distinct BicD homologs with distinct coiled-coil registries have been determined. *Hs* BicD2-CTD has a homotypic coiled-coil registry, with characteristic layers of knobs-into-holes interactions, which are formed by the same knob residues from both chains, and therefore the helices are aligned at equal height. Wild-type *Dm* BicD-CTD (Fig. 2A) however has an asymmetric coiled-coil registry. The N-terminal half of the coiled-coil has a heterotypic registry, in which residue *i* from one chain is paired up with residue *i*+4 from the second chain to form layers of knobs-into-holes interactions, resulting in a vertical displacement of the helices by ∼one helical turn. In the C-terminal half, the helices are aligned to form a coiled-coil with homotypic registry.

To determine the coiled-coil registry, we assigned the heptad register of the structure of *Dm* BicD-CTD/F684I (Fig. 2, Fig. S1). As observed for the wild-type *Dm* BicD-CTD (Fig. 2A) and *Hs* BicD2-CTD (Fig. 2C), the C-terminal half of the mutant structure has a homotypic coiled-coil registry (Fig. 2B, see Table 2 for residue numbering in *Hs* and *Dm* homologs). Notably, one additional layer of knobs-into-holes interactions with homotypic coiled-coil registry was identified in the Bicaudal mutant compared to the *Dm* BicD wild type. This layer is formed by knob residue V702 from both chains (Fig. 2, Fig. S1). In contrast to the F684I mutant, in the *Dm* BicD wild type, V702 forms a layer of knobs-into-holes interactions with residue Y698 and therefore part of the region that has a heterotypic coiled-coil registry. Wild-type *Dm* BicD-CTD also has additional layers of knob-into-holes interactions with heterotypic registry in its N-terminal region, resulting in a vertical displacement of the helices by approximately one helical turn. However, no additional layers of knobs-into-holes interactions N-terminal of V702 were identified in the structure of *Dm* BicD-CTD/F684I, because a ∼20 residue region upstream of residue I684 is not resolved for one chain (Fig. 2).

**Table 2.** Numbering of homologous key residues in *Dm* BicD and *Hs* BicD2.

|  | ***Dm* BicD** | ***Hs* BicD2** |
| --- | --- | --- |
| Conversion | Residue *i* | Residue *i*+59 |
| C-terminal domain (CTD) | 656-745 | 715-804 |
| Bicaudal mutant | F684I | F743I |
| Key aromatic residues | F684, F691, Y698 | F743, F750, Y757 |
| Nup358 interface residues* | L687, R688, M690, R694 | L746, R747, M749, R753 |

\*Interface residues were mapped for *Ms* BicD1, a close homolog of *Hs* BicD2 (Terawaki *et al.*, 2015). Residue *i* of *Hs* BicD2 (as listed in the table) is homologous to residue *i*-2 of *Ms* BicD1.

A least-squares superimposition of the structures of the *Dm* BicD-CTD/F684I mutant and the wild type revealed that in the mutant, both I684 residues are aligned at equal height, as observed for the homologous residue F743 in *Hs* BicD2 with homotypic registry (Fig. 2 D-G). However, in the wild-type *Dm* BicD structure, which has an asymmetric coiled-coil registry, residues F684 from both chains are vertically shifted by approximately one helical turn with respect to each other (Fig. 2 D-G). F684 from one chain lines up with R688 from the second chain to form a layer of knobs-into-holes interactions (heterotypic registry) (Fig. 2A).

To conclude, several pieces of data suggest that the structure of the *Dm* BicD2-CTD/F684I mutant has a homotypic coiled-coil registry: One additional layer of knobs-into-holes interactions with homotypic coiled-coil registry is formed by residues V702 in the structure of the Bicaudal mutant compared to the wild type. Furthermore, residues I684 are aligned in the structure of the Bicaudal mutant at equal height (homotypic registry), whereas in the wild-type structure, residues F684 are vertically shifted respectively to each other by approximately one helical turn (heterotypic registry). However, since a ∼20 residue area of one monomer in the Bicaudal mutant is not resolved, no additional knobs-into-holes interactions were identified, therefore, the coiled-coil registry N-terminal of I684 is unknown.

### Distinct conformations of F684 and F691 stabilize distinct coiled-coil registries

Another key residue that is important to stabilize either a homotypic or asymmetric coiled-coil registry is F691 (Fig. 3). In the *Dm* BicD-CTD wild-type structure with the asymmetric coiled-coil registry, phenylalanine side chains of F691 from both chains interact in a face-to-edge aromatic interaction, which leads to vertical displacement of the chains by ∼1 helical turn (Fig. 3B). However, in the BicD homologs with homotypic registry (*Ms* BicD1, *Hs* BicD2) the homologous phenylalanine residues interact with face-to-face aromatic interaction, which allows the chains to be aligned in the homotypic registry (Fig. 3D, E). In the *Dm* BicD F684I mutant, the phenylalanine side chains of F691 from both chains interact face-to-face (Fig. 3A, C, F), which suggests a homotypic registry. A least-squares superimposition of the structures of the *Dm* BicD wild type and the F684I mutant confirms that the F691 side chains form a face-to-face aromatic interaction in the structure of the Bicaudal mutant and a face-to-edge aromatic interaction in the wild-type structure with the asymmetric registry (Fig. 3C). These data suggest that in the structure of the *Dm* BicD F684I mutant, the F691 residues assume a conformation that is found in *Hs* BicD2 with homotypic coiled-coil registry.

**Figure 3.**
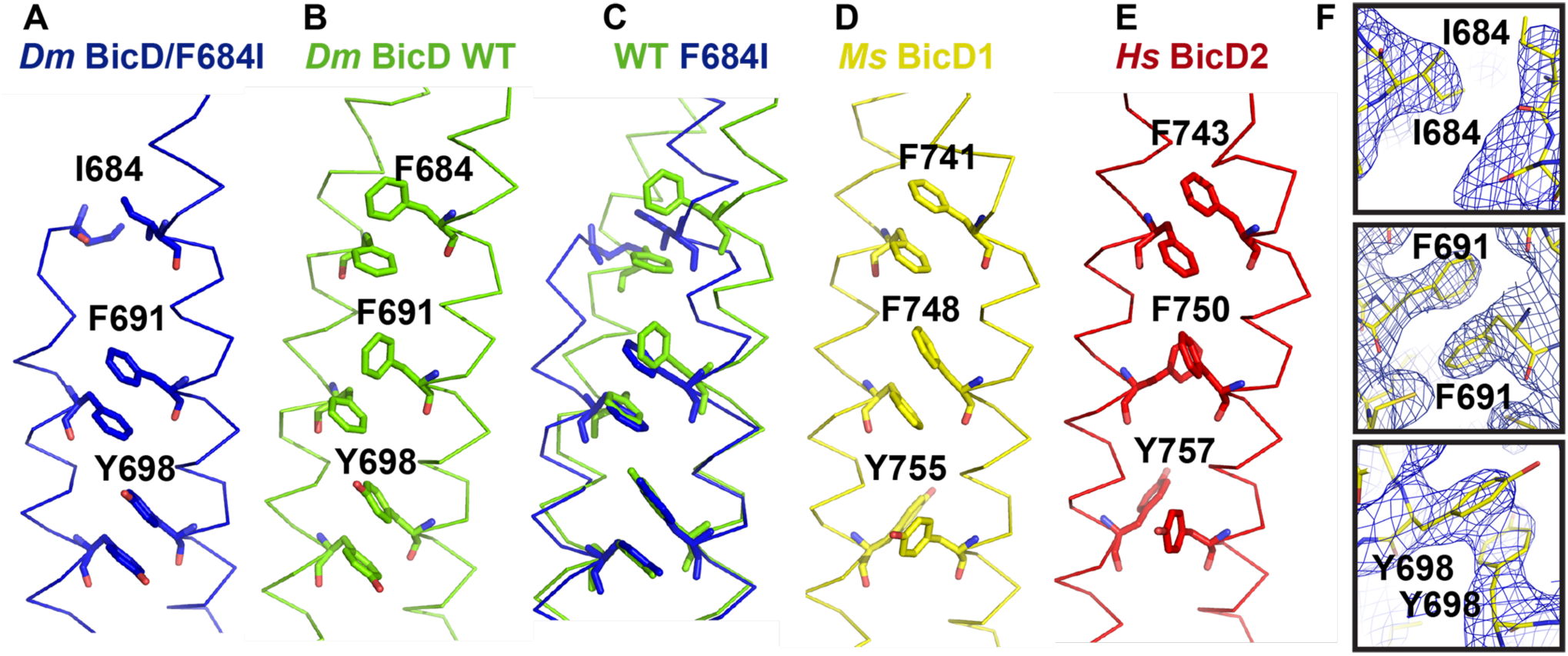
Conformation of key aromatic residues in *Dm* BicD-CTD/F684I. (A) The Cα-trace of the structure of *Dm* BicD-CTD/F684I is shown (blue). Residues I684, F691, Y698 are labeled and shown in stick representation. (B) Structure of the wild-type *Dm* BicD-CTD (green, PDB ID 4BL6) (Liu *et al.*, 2013). (C) Least squares superimposed structures of *Dm* BicD-CTD/F684I and the wild type. (D, E) Structures of the *Dm* BicD homologs (D) *Ms* BicD1-CTD (yellow, PDB ID 4YTD) (Terawaki *et al.*, 2015) and (E) *Hs* BicD2-CTD (red, PDB ID 6OFP) (Noell *et al.*, 2019). (F) Structure of the *Dm* BicD-CTD/F684I mutant in stick representation overlaid with the 2F_O_-F_C_ electron density map (blue mesh). Close-ups of residues I684, F691 and Y698 are shown in three panels. Note that residues F691 from both chains are oriented face-to-face, as observed in the structures with the homotypic registries.

### The disordered region is present in the crystal, and α -helical

Because a ∼20 residue region of one chain is not resolved in the crystal structure, we dissolved crystals of *Dm* BicD-CTD/F684I and analyzed them by SDS-PAGE, to assess whether the crystals contained the intact protein (Fig. 4A). A comparison of the SDS-PAGE of the dissolved crystals, the purified *Dm* BicD-CTD/F684I protein as well as the wild-type protein suggests that indeed the intact *Dm* BicD-CTD/F684I protein is present in crystal, suggesting that the unresolved N-terminal region is disordered.

**Figure 4.**
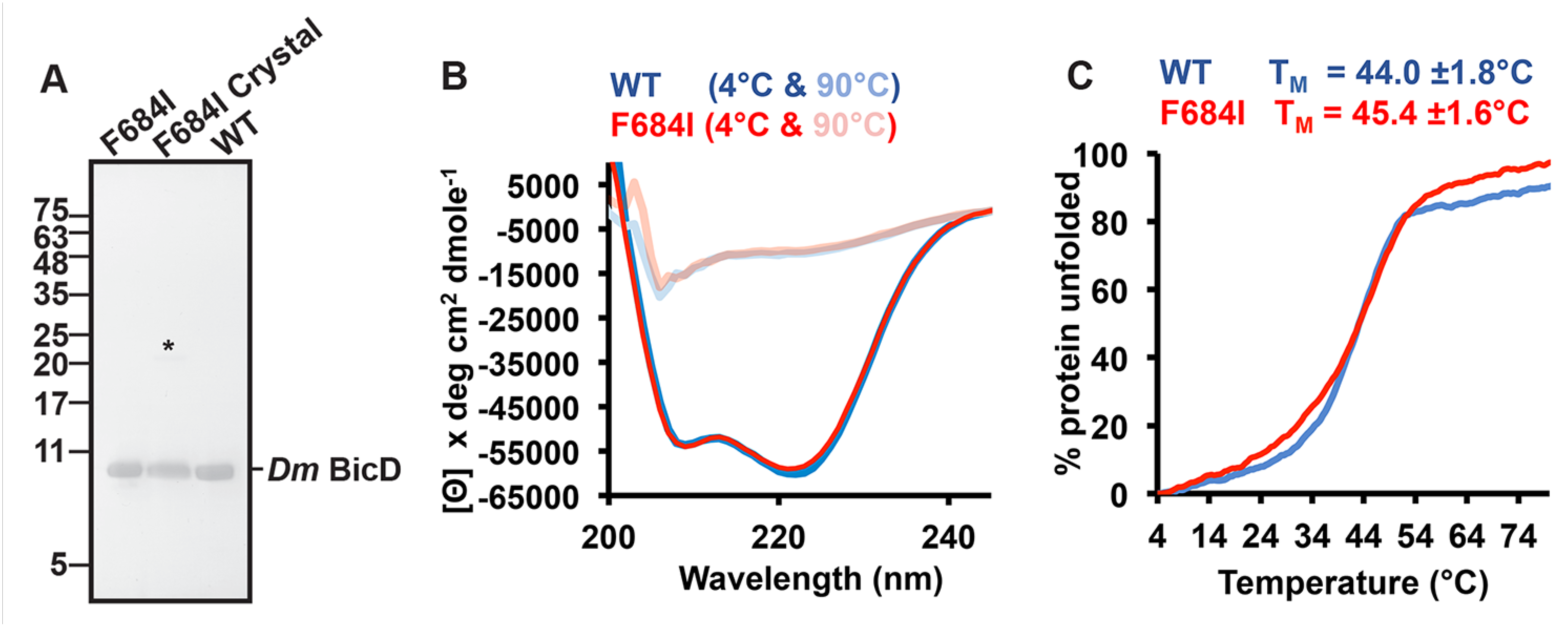
The intact *Dm* BicD-CTD/F684I is present in the crystal and it is fully folded. (A) SDS-PAGE analysis of purified *Dm* BicD-CTD/F684I (left lane), *Dm* BicD-CTD/F684I crystals (middle lane), and purified wild-type protein (right lane). The position of the dimer band is indicated by an asterisk. For the crystal sample, 20 crystals were washed three times with reservoir buffer before being dissolved in gel filtration buffer. (B) CD wavelength scans of *Dm* BicD-CTD WT (blue) and *Dm* BicD-CTD/F684I (red) at 4°C (native) and 90°C (random coil). The mean residue molar ellipticity [Θ] versus the wavelength is shown. Experiments were repeated three times, representative scans are shown. See also Fig. S2. (C) Thermal unfolding curves of wild type (blue) and F684I (red) were recorded by CD spectroscopy at 222 nm. Molar ellipticity [Θ] versus temperature is plotted. 0% and 100% protein unfolded represent the values of [Θ]_min_ and [Θ]_max_, respectively. Representative experiments are shown; melting temperatures T_M_ of *Dm* BicD-CTD WT and F684I are shown and were averaged from three experiments.

In order to assess, whether the disordered portion of the helix is α-helical (rather than misfolded), we probed the secondary structure content of *Dm* BicD-CTD/F684I by circular dichroism (CD) wavelength scans. The CD wavelength spectra of the *Dm* BicD-CTD/F684I mutant and the wild-type both have minima at 208nm and 222nm, which are characteristic for α-helical proteins. Notably, the spectra of the F684I mutant and the wild-type overlay perfectly, suggesting that both structures have a very similar α-helical content. These data suggest that the ∼20 residue disordered region is α-helical rather than unstructured (Fig. 4B). In order to assess, whether differences in the crystallization conditions of the *Dm* BicD-CTD/F684I mutant and the wild-type protein contributed to the observed structural differences, we also recorded CD wavelength spectra in modified crystallization buffers (Fig. S2). These spectra confirmed that the compounds of the crystallization buffers do not affect the α-helical content of either the mutant or the wild-type protein, and therefore do not cause the observed structural disorder in the mutant.

Furthermore, we compared the dimer interface of the structure of the F684I mutant with the wild type (Table S1). Since the N-terminal region of one of two chains is disordered in the mutant, the interface area is smaller (1427 Å^2^) compared to the wild type (1764 Å^2^), and the dimer interface of the F684I mutant contains eighteen fewer interacting residues as well as one less hydrogen bond and one less salt bridge compared to the wild type (Table S1). It is unknown if the disordered region engages in interactions that stabilize the dimer, however, in the absence of additional interactions one would expect that the F684I mutant would be less stable than the wild type.

Thus, we probed thermodynamic stability of the F684I mutant and the wild type by recording circular dichroism spectroscopy melting curves (Fig. 4C). Protein unfolding was monitored by CD spectroscopy at 222 nm. The apparent melting temperature T_M_ of *Dm* BicD-CTD/F684I was 45.4±1.6°C, which is similar to the T_M_ of the wild-type protein (44.0±1.8°C) (Fig. 4C). Based on the melting temperatures, the *Dm* BicD-CTD/F684I mutant has comparable thermodynamic stability as the wild type, despite the N-terminal disordered portion of one helix. Therefore, it is conceivable that the disordered portion still interacts with the other chain, as it would explain the observed similar thermodynamic stability.

To conclude, the ∼20 residue disordered region is present in the crystal and folded, and since the thermodynamic stability of the mutant is comparable to the wild type, this region is likely to still interact with the other chain. These data suggest that the region N-terminal of I684 is flexible in the mutant in one chain and possibly assuming multiple conformations.

### MD simulations suggest that the N-terminal region of the mutant can switch between homotypic and heterotypic registries

In the X-ray structure, the Bicaudal mutant of *Dm* BicD-CTD assumes a conformation with a homotypic coiled-coil registry, and the region N-terminal of F684I is disordered for one of two chains. In order to gain insight into the disorder in the N-terminal region of *Dm* BicD-CTD/F684I, we used MD simulations to assess if a conformation of the mutant with a homotypic registry would sample multiple conformations. For these simulations, the structure of the homolog *Hs* BicD2-CTD was chosen as a starting point, since it has a fully resolved homotypic coiled-coil registry (unlike *Dm* BicD WT), and amino acid mutations were carried out to match the sequence of *Dm* BicD-CTD/F684I. In these simulations, the N-terminal region of the coiled-coil switched back and forth between a homotypic and heterotypic registry, while the C-terminal region retained a homotypic registry, in line with the various crystal structures. Therefore, the overall coiled-coil registry of *Dm* BicD-CTD/F684I switched after ∼53 ns from homotypic to asymmetric, and reverted back to homotypic after ∼120 ns (Fig. 5A-C). This suggests that the disorder in the N-terminal region of one of the chains, as seen in the crystal structure, is likely caused by the ability of the N-terminal region to easily switch between the homotypic and heterotypic registries.

**Figure 5.**
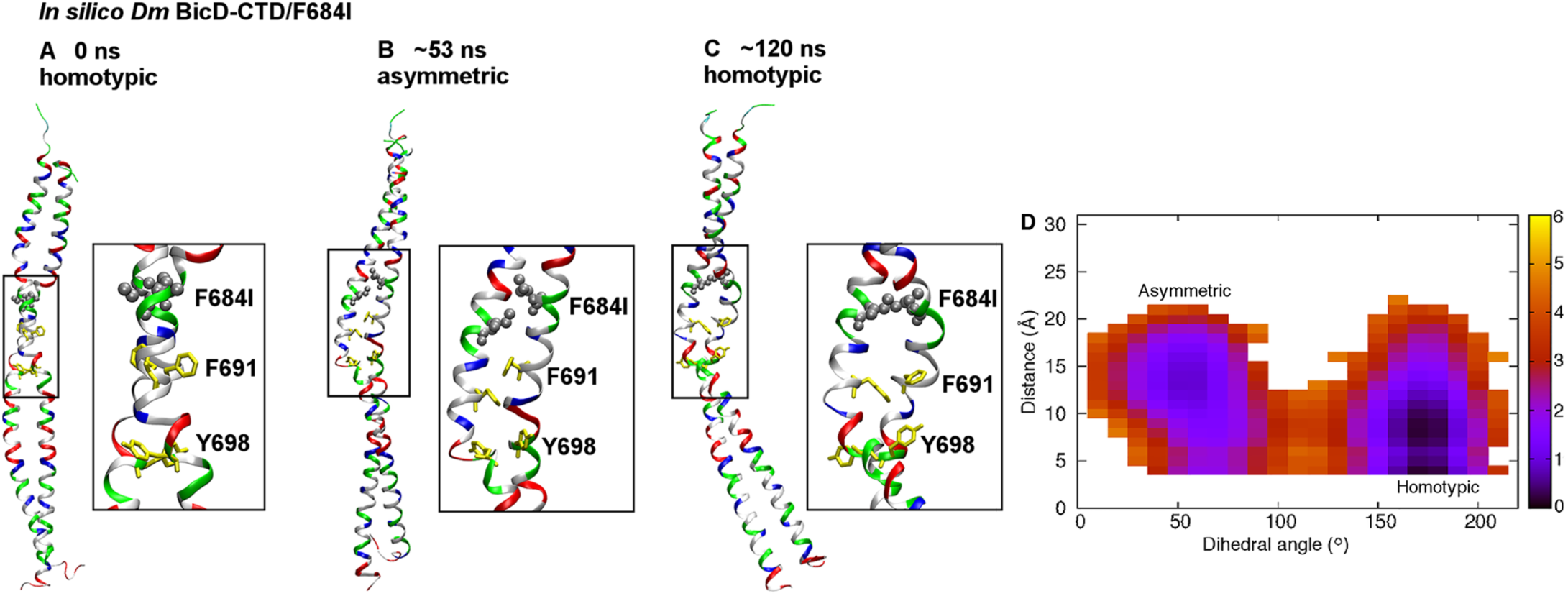
MD simulations suggest that the *Dm* BicD-CTD/F684I mutant switches between homotypic and asymmetric registries, with the N-terminal region switching between homotypic and heterotypic registries. (A) Cartoon representation of the equilibrated structure of *Dm* BicD-CTD/F684I (Noell *et al.*, 2019), with homotypic registry, colored by residue type (blue: positively charged, red: negatively charged, green: polar, white: non-polar). F684 was mutated to isoleucine (silver spheres). F691 and Y698 are shown in yellow stick representation. A close-up of the boxed area is shown on the right. See also File S1. (B) Structure of the F684I mutant of *Dm* BicD-CTD after ∼53 ns of an MD simulation. Note that the N-terminal region of the coiled-coil switches to a heterotypic registry; therefore, the overall coiled-coil registry is asymmetric. See also File S2. (C) Structure of the F684I mutant of *Dm* BicD-CTD after ∼120 ns of the same MD simulation. Note that the structure switches back to a homotypic coiled-coil registry. However, the solvent-exposed F691 sidechains are oriented towards the same side, as opposed to opposite sides in A. This leads to a slight distortion of the coiled coil around the F691 residues. See also File S3. (D) Free energy in kcal/mol as a function of the C-C_α_-C_β_-C_γ_ dihedral angle of F691 of chain A (plotted along the horizontal axis), and the distance between the sidechain N atom of K678 of chain A and the C_δ_ atom of E673 of chain B (plotted along the vertical axis). The distance between the sidechain N atom and C_δ_ was chosen, since both oxygen atoms of the carboxyl group can engage in salt bridge formation. The free energy is depicted using a color map that ranges from 0 to 6 kcal/mol. The free energy difference between the minima is ∼1 kcal/mol, with a free energy barrier of ∼4-5 kcal/mol.

In order to gain insights into the kinetics of the observed coiled-coil registry shift, we also calculated the relative free energies of the conformations with homotypic and asymmetric coiled-coiled registries, as well as the free energy barrier that separates them. Detailed analysis of the MD trajectory for *Dm* BicD-CTD/F684I revealed the C-C_α_-C_β_-C_γ_ dihedral angle of F691 of chain A to be directly correlated with the registry shift. It assumed values around 175° and around 55° in conformations with homotypic and asymmetric registries, respectively. Interestingly, the corresponding dihedral angle of F691 of chain B and of I684 and Y698 of either chain were not found to be correlated with the registry shift. The different behaviors of F691 of the two chains is consistent with the disorder in the N-terminal region of only one of the chains in the crystal structure. In addition to the above mentioned dihedral angle, the salt-bridge interaction between K678 of chain A and E673 of chain B that is formed in the conformation with the homotypic registry was also found to be correlated with the registry shift, with the interaction completely broken in the conformation with asymmetric registry. Interestingly, of all the salt-bridges in the N-terminal region, only this one was found to be related to the registry shift. Our data therefore provide insights into the molecular mechanism of the coiled-coil registry shift, and reveal key roles of residue F691 from chain A as well as of the salt bridge between K678 of chain A and E673 of chain B in the mechanism.

The identification of these key coordinates related to registry shift allowed the calculation of the potential of mean force (PMF) or free energy as a function of the two coordinates (Eqn. 1, see Materials and Methods), revealing the free energy difference between the homotypic and asymmetric registries to be less than 1 kcal/mol, and the free energy barrier for transition between the registries to be ∼4-5 kcal/mol (Fig. 5D). Hence, the two registries have similar stability and can interconvert on a timescale of tens of ns, as observed in the MD simulations. In the crystal structure of *Dm* BicD-CTD WT, disorder was not observed. It is conceivable that F684 is a key residue that serves to lock the WT BicD coiled-coil in conformations with either homotypic or asymmetric registry. Replacement of this residue by isoleucine may lead to promiscuity, allowing other conformations to form. This is in line with the crystal structure of the F684I mutant and the MD simulations. It should be noted that the structure of the mutant has an elevated overall B-factor (Table 1), indicating flexibility, and a region of one chain that undergoes coiled-coil registry shifts is also disordered, further suggesting conformational variability.

In comparison, in our recent MD simulations of the human homolog of the Bicaudal mutant, *Hs* BicD2-CTD/F743I with asymmetric coiled-coil registry, the homologous F743I mutation induced a coiled-coil registry shift from an asymmetric to a fully heterotypic registry (Noell *et al.*, 2019). Simulations starting from the homotypic registry of *Hs* BicD2-CTD/F743I maintained a homotypic registry (Figure S3), suggesting that the human homolog of the Bicaudal mutant can also sample multiple registries. Hence conformational flexibility caused by the F684I mutation is not unique to *Dm* BicD-CTD, but also present in other homologs.

To conclude, our MD simulations of the Bicaudal mutant show that it samples conformations with homotypic and asymmetric coiled-coil registries which are of similar stability on a time scale of tens of ns (Fig. 5), which would explain why one chain in the N-terminal region of the coiled-coil is disordered in the crystal structure.

### The homologous F743I mutation modulates cargo selection in Hs BicD2

Because a registry shift is expected to re-model the surface of the coiled-coil, which harbors binding sites for cargo adaptors, we investigated whether a coiled-coil registry shift in BicD2 plays a role in cargo selection. The binding sites for the cargo adaptors *Dm* Egalitarian and *Dm* Rab6^GTP^ on *Dm* BicD have been previously mapped to residues 702-743 (Fig. 6A) (Liu *et al.*, 2013; Terawaki *et al.*, 2015). A similar region has been mapped as minimal Rab6^GTP^ binding site for a close homolog of human BicD2 (residues 755-802, Fig. 6B, see Table 2 for residue numbering in *Dm* and *Hs* homologs). These cargo adaptors bind to the C-terminal homotypic region of *Dm* BicD-CTD/ *Hs* BicD2-CTD, which does not undergo coiled-coil registry shifts. Therefore the F684I mutation, which likely induces a coiled-coil registry shift, and which is located N-terminal of the mapped binding sites, is not expected to alter cargo adaptor binding. Indeed, the F684I mutant of *Drosophila* BicD does not affect the interaction of BicD with the cargo adaptors Egalitarian and Rab6^GTP^ (Liu *et al.*, 2013).

**Figure 6.**
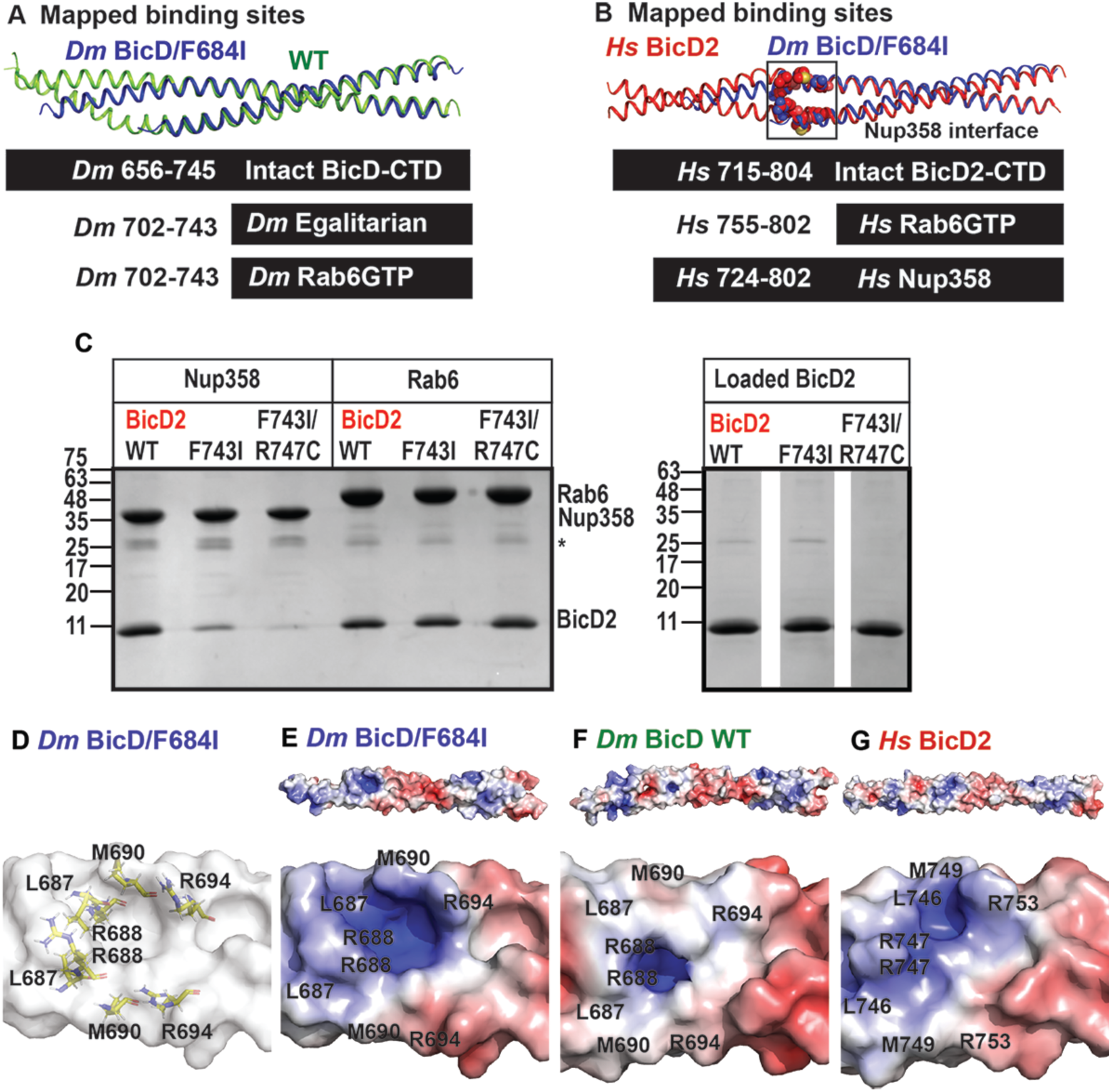
Role of coiled-coil registry shifts in cargo selection of human BicD2. (A) Least-squares superimposition of the structures of *Dm* BicD-CTD/F684I (dark blue) and wild-type (green) in cartoon representation is shown. A schematic representation of the intact *Dm* BicD-CTD (black bar) and of the mapped cargo adaptor binding sites including residue numbers is shown below. (B) Least squares superimposition of the structures of *Dm* BicD-CTD/F684I (dark blue) and *Hs* BicD2-CTD (red) in cartoon representation, with a schematic representation of the intact protein and the mapped cargo adaptor binding sites below. Known Nup358/BicD2 interface residues (see Table 2) are shown in spheres representation (Terawaki *et al.*, 2015). (C) Pull-down-assays of BicD2-CTD (wild type, F743I or F743I/R747C mutant) with the GST-tagged cargo adaptors Rab6^GTP^-GST and Nup358-min-GST. An asterisk indicates the location of the GST-band. An SDS-PAGE of the elution fractions is shown in the left panel. Right panel: SDS-PAGE analysis of the BicD2-CTD load fractions. Pull-down assays were repeated three times with similar results. (D) Surface representation of the structure of *Dm* BicD-CTD/F684I. Several important Nup358/BicD2 interface residues are known (Terawaki *et al.*, 2015) (see Table 2); homologous *Dm* residues are shown in stick representation and labeled. (E-G) The surface electrostatic potential of distinct structures is shown. Positive (blue: 5 kT/e) and negative (red: −5 kT/e) potentials are mapped on the solvent excluded molecular surface (top panel). The bottom panel shows a close-up of the known BicD2/Nup358 interface residues. The same view as in (D) is shown. (E) *Dm* BicD-CTD/F684I. (F) *Dm* BicD-CTD wild type (Liu *et al.*, 2013). (G) *Hs* BicD2-CTD (Noell *et al.*, 2019).

Notably, the mammalian cargo adaptor Nup358 binds to a larger region that includes residues 724-802 of *Hs* BicD2 (mapped for a homolog) (Terawaki *et al.*, 2015), and a portion of this region undergoes a coiled-coil registry shift (Fig. 6B). Several important Nup358 interface residues, which are N-terminal of F743 were identified for a close homolog of human BicD2 (Table 2; residues L746, R747, M749 and R753). Mutation of each of these residues to alanine strongly diminishes the interaction (Terawaki *et al.*, 2015).

We therefore hypothesized that the homologous F743I mutation in *Hs* BicD2 would modulate the interaction between BicD2 and Nup358, since a portion of the interface is located in the region that is thought to undergo coiled-coil registry shifts. The BicD2/Rab6^GTP^ interaction however is expected to be unaffected by the F743I mutation, since the binding site is located in the region that remains homotypic. This is confirmed by pull-down assays of Rab6^GTP^ with the BicD2-CTD/F743I mutant, and the F743I/R747C mutant, which both bind with comparable strength as observed for the wild-type (Fig. 6C). Interestingly, binding of Nup358 is modulated by the mutation. In a pull-down-assay, the minimal interacting domain Nup358-min pulls down wild-type BicD2-CTD much more strongly compared to the F743I mutant (Fig. 6C). Binding is even more strongly weakened for the double mutant F743I/R747C, which combines two mutations that induced coiled-coil registry shifts in simulations (Fig. 6C) (Noell *et al.*, 2019).

To gain further insights into the impact of the mutation on the cargo-adaptor-binding interface, we compared the electrostatic surface potential of the structures of *Dm* BicD-CTD/F684I, the wild type (asymmetric registry), and *Hs* BicD2-CTD (homotypic registry) (Fig. 6D-G). There are differences in the electrostatic surface potential in the area where Nup358 binds (L746, R747, M749 and R753 (Terawaki *et al.*, 2015) (Fig. 6D-G). Notably, both the Bicaudal mutant BicD-CTD/F684I and *Hs* BicD2-CTD (homotypic registry) have a highly positively charged surface electrostatic potential in the area of these interface residues, creating a basic pocket (blue, Fig. 6E,G). In comparison, the same interface area in the *Dm* BicD-CTD has a much less charged electrostatic surface potential (Fig. 6F). Such changes could be caused by a coiled-coil registry shift and could be responsible for the observed difference for the interaction between *Hs* BicD2-CTD wild type and the F743I mutant with Nup358-min.

To conclude, while the *Dm* BicD/F684I mutant shows comparable selectivity as the wild-type protein towards the cargo adaptors Egalitarian and Rab6^GTP^ (Liu *et al.*, 2013), in human BicD2, the homologous F743I mutant, which likely induces a coiled-coil registry shift, affects cargo selection. While binding of BicD2 to Rab6^GTP^ is not affected by the mutation, binding to Nup358 is strongly reduced, likely because it binds to a larger binding site that contains a portion of the protein that may undergo a coiled-coil registry shift.

## DISCUSSION

In the absence of cargo, BicD forms an auto-inhibited state that is unable to recruit dynein (Chowdhury *et al.*, 2015; Liu *et al.*, 2013; McClintock *et al.*, 2018; McKenney *et al.*, 2014; Schlager, Hoang, *et al.*, 2014; Schlager, Serra-Marques, *et al.*, 2014; Sladewski *et al.*, 2018; Splinter *et al.*, 2012; Terawaki *et al.*, 2015; Urnavicius *et al.*, 2015). Here we show by single molecule processivity assays that auto-inhibition is abolished in the classical Bicaudal mutant BicD/F684I, resulting in cargo-independent activation of dynein-dynactin, consistent with cellular studies that show increased dynein recruitment (Liu *et al.*, 2013; Mohler & Wieschaus, 1986). To probe the mechanism of BicD activation from the auto-inhibited state, we determined the X-ray structure of the C-terminal cargo-binding domain of the Bicaudal mutant (*Dm* BicD-CTD/F684I). The Bicaudal mutant assumes a conformation with homotypic registry as its predominant structural state, as the helices are aligned at equal height up to residue I684, unlike in the wild-type, where F684 residues are displaced vertically by one helical turn against each other. However, a ∼20 residue region upstream of residue I684 is not resolved for one chain in the structure. This structural flexibility is also confirmed by MD simulations and free energy calculations, in which the mutant samples conformations with homotypic and asymmetric coiled-coil registries of similar stability on a time scale of tens of ns, which would explain the observed disorder in the crystal structure. Our data suggest that the F684I mutation shifts the equilibrium of registry-shifted conformers, resulting in formation of a larger percentage of BicD with homotypic registry.

It was previously proposed that BicD undergoes coiled-coil registry shifts, which activate it for dynein recruitment upon cargo binding (Liu *et al.*, 2013; Noell *et al.*, 2019; Terawaki *et al.*, 2015). This idea is based on the structures of distinct BicD homologs with distinct coiled-coil registries as well as our recent MD simulations, which suggest that human BicD2 can assume stable conformations with either homotypic or asymmetric coiled-coil registries (Liu *et al.*, 2013; Noell *et al.*, 2019; Terawaki *et al.*, 2015).

Here, we show that in MD simulations, the structure of the *Dm* BicD/F684I mutant transitions back and forth between homotypic and asymmetric coiled-coil registries on a time scale of tens of ns. These simulations also reveal key roles of residue F691 from chain A as well as for the salt bridge between K678 of chain A and E673 of chain B in the structural transition. Our results suggest that the mutation induces structural dynamicity, and leads to formation of multiple conformations, which is also supported by the crystal structure. The structure has a comparatively high B-factor, suggesting flexibility and a ∼20 amino acid region at the N-terminus of one chain is not resolved in the structure, whereas the second chain is resolved, and therefore the coiled-coil registry in this region cannot be determined. It is unlikely that the different crystallization conditions contribute to the distinct conformations of the coiled-coil, since the mutant and wild-type protein have the same α-helical content in different crystallization buffers. Our data suggest that the disordered region is present in the crystal and α-helical. The disordered region likely also still interacts with the second ordered chain, since the melting temperatures in solution studies indicate similar thermodynamic stability for the mutant and the wild-type.

In the wild-type, F684 likely serves as a switch to lock *Dm* BicD in conformations with distinct registries (Liu *et al.*, 2013; Noell *et al.*, 2019). In the conformation with the asymmetric registry, F684 rotates to the core of the coiled-coil and forms an edge-to-face aromatic interaction with residue F684 from the second chain, which leads to the observed vertical displacement of the helices (Liu *et al.*, 2013; Noell *et al.*, 2019). In the mutant, the phenylalanine side chain is replaced by a much smaller isoleucine side chain, which likely cannot lock the asymmetric registry in place, and due to its smaller size is unable to prevent conformational changes. The result is likely a dynamic mixture of several states.

The single molecule assays, which use full-length mutant BicD, are consistent with the predictions derived from the minimal cargo-binding domain structure. In the auto-inhibited conformation, full-length BicD2 forms a looped structure in which the CTD binds to the N-terminal dynein/dynactin binding site, likely causing steric interference (Chowdhury *et al.*, 2015; Liu *et al.*, 2013; McClintock *et al.*, 2018; McKenney *et al.*, 2014; Schlager, Hoang, *et al.*, 2014; Schlager, Serra-Marques, *et al.*, 2014; Sladewski *et al.*, 2018; Splinter *et al.*, 2012; Terawaki *et al.*, 2015; Urnavicius *et al.*, 2015). Cargo binding could induce a local coiled-coil registry shift in the BicD2-CTD, which might be sufficient to weaken binding to the N-terminal dynein/dynactin binding site and thus activate BicD for dynein recruitment. Alternatively, the registry shift could propagate through the entire coiled-coil to the N-terminal dynein-dynactin binding site. Furthermore, the induced flexibility and formation of multiple conformations as observed in the mutant may potentially also be an inherent structural feature of cargo-bound wild type BicD. Such structural and mechanistic details remain to be established and studies with full-length proteins in physiological context remain to be conducted to fully understand the molecular mechanism of BicD2 auto-inhibition and activation.

In addition to BicD2, several other dynein adaptors have coiled-coil structures (e.g. NudE/NudEL (Efimov & Morris, 2000; Niethammer *et al.*, 2000; Stehman *et al.*, 2007), the Hook proteins (Bielska *et al.*, 2014; Olenick *et al.*, 2019), RILP (Cantalupo *et al.*, 2001), Rab11-FIP3 and Spindly (Griffis *et al.*, 2007; Mosalaganti *et al.*, 2017) and some of them, e.g. Spindly assume an auto-inhibited state in the absence of cargo (McKenney *et al.*, 2014; Mosalaganti *et al.*, 2017). The molecular mechanism for auto-inhibition in these dynein adaptors remains to be established, and it is possible that some of them undergo coiled-coil registry shifts as well.

Cargo selection for BicD2-dependent transport events are tightly regulated, but currently known regulatory mechanisms, which include competition of cargo adaptors (Noell *et al.*, 2018) and the G2 specific kinase Cdk1 (Baffet *et al.*, 2015), are insufficient to fully explain how BicD2 switches between these cargoes in a cell-cycle specific manner (Noell *et al.*, 2018).

In *Drosophila*, cargo adaptors Egalitarian and Rab6^GTP^ bind to a small domain of BicD that remains homotypic and does not undergo coiled-coil registry shifts, consistent with the Bicaudal mutation not affecting cargo selection (Dienstbier *et al.*, 2009; Liu *et al.*, 2013). The F684I mutation does not affect the affinity of BicD for Egalitarian or Rab6^GTP^ (Liu *et al.*, 2013) but it promotes cargo-independent dynein recruitment (Fig. 1) thereby resulting in increased transport rates. An increase in dynein-mediated transport of *Oskar* mRNA/Egalitarian in the F684I mutant causes the double-abdomen fly phenotype (Bull, 1966; Liu *et al.*, 2013; Mach & Lehmann, 1997; Mohler & Wieschaus, 1986; Navarro *et al.*, 2004; Zimyanin *et al.*, 2008). This likely means that dynein recruitment is more limiting to transport than the affinity of BicD towards cargo adaptors (Bullock *et al.*, 2006; Liu *et al.*, 2013).

Notably, we propose that coiled-coil registry shifts in human BicD2 modulate cargo selection. Human Nup358 binds to a larger interface on BicD2 that includes a region which undergoes coiled-coil registry shift (Terawaki *et al.*, 2015). The homologous F743I mutation which is expected to induce a coiled-coil registry shift (Noell *et al.*, 2019) diminishes binding of Nup358 to *Hs* BicD2, whereas the interaction of BicD2 with Rab6^GTP^ is unaffected. Binding of cargo-adaptors such as Nup358 is expected to induce a coiled-coil registry shift in BicD2, and in our binding assays, Nup358 remains associated with BicD2. It remains to be established by future studies, if this is due to a slow dissociation constant of the complex or whether the interaction with BicD2 induces structural changes in Nup358 that stabilize it in the complex once bound (“induced fit”). Notably, a registry shift could have a regulatory role in preventing the cargo adaptor Nup358 from binding. A coiled-coil registry shift in BicD2 could be triggered by a regulatory signal, which could modulate cargo selection by reducing the binding of selected cargo adaptors including Nup358.

In conclusion, our data provide mechanistic insights into auto-inhibition and cargo selection of the dynein adaptor BicD2. Our results suggest that the full-length *Dm* BicD/F684I mutant is capable of activating dynein for processive transport in a cargo-independent manner. The X-ray structure of *Dm* BicD-CTD/F684I reveals that the mutation induces a coiled-coil registry shift, which we propose as the underlying mechanism for cargo-independent activation. A ∼20 residue N-terminal region of one monomer is disordered in the structure, in line with MD simulations of the mutant which samples conformations with homotypic and asymmetric registries on a time scale of tens of ns. Free energy calculations indicate that conformations with homotypic and asymmetric registries have similar stability and are separated by a free energy barrier of ∼4-5 kcal/mol. The observed structural dynamicity could either be a structural feature of activated BicD2 or it could be caused by the mutation. Notably, the human homolog of the Bicaudal mutant shows reduced affinity to the cargo adaptor Nup358, which recruits BicD2 to the nuclear envelope. We thus propose that a coiled-coil registry shift modulates cargo selection for BicD2-dependent transport pathways, which are important for cell cycle control and brain development (Baffet *et al.*, 2015; Bianco *et al.*, 2010; Dienstbier *et al.*, 2009; Hu *et al.*, 2013; Matanis *et al.*, 2002; Splinter *et al.*, 2010).

## MATERIALS AND METHODS

### Protein expression and purification

Codon optimized human dynein for expression in Sf9 cells (DYNC1H1 (DHC), DYNC1I2 (DIC), DYNC1LI2 (DLIC), DYNLT1 (Tctex), DYNLRB1 (Robl) and DYNLL1(LC8)) was a generous gift from Simon Bullock (Schlager, Hoang, *et al.*, 2014). The heavy chain was modified to contain an N-terminal FLAG tag followed by a biotin tag to enable heavy chain labeling. Dynein expression in Sf9 cells and purification was as described in (Sladewski *et al.*, 2018). Purified dynein was stored at −20°C in 10 mM imidazole, pH 7.4, 0.2 M NaCl, 1 mM EGTA, 2 mM DTT, 10 µM MgATP, 5 µg/ml leupeptin, 50% glycerol. Dynactin was purified from ∼300 g bovine brain tissue as described (Bingham *et al.*, 1998), and stored at −20°C in the same buffer as dynein. Full-length WT and mutant *Drosophila* BicD (BicD^WT^ and BicD^F684I^) were expressed in Sf9 cells and *Drosophila* BicD-coiled-coil domain 1 (BicD^CC1^) in bacteria as described (11).

To create an expression construct of *Drosophila* BicD-CTD/F684I (residues 656-745), the corresponding codon optimized DNA sequence was commercially synthesized by Genscript and cloned into the pET28a vector with the NdeI and XhoI restriction sites. *Drosophila* Bicaudal D-CTD F684I (*Dm* BicD-CTD F684I, residues 656-745) was expressed in E. *coli* BL20(DE3)-RIL strain. His_6_-tagged *Dm* BicD-CTD/F684I was purified by Ni-NTA affinity chromatography and the tag was cleaved by thrombin, followed by second Ni-NTA affinity chromatography as described in (Noell *et al.*, 2018). The protein was further purified by gel filtration chromatography on a HiLoad™ 16/600 Superdex 200 pg column (GE Healthcare) with the following buffer: 20 mM HEPES pH 7.5, 150 mM NaCl, 0.5 mM TCEP as described (Noell *et al.*, 2018). Wild-type *Dm* BicD-CTD was expressed and purified as described (Cui *et al.*, 2018; Loftus *et al.*, 2017; Noell *et al.*, 2019). Purified protein was analyzed by SDS-PAGE, 16% acrylamide gels and stained by Coomassie Blue.

GST-pull down assays of Rab6^GTP^-GST, Nup358-min-GST and *Hs* BicD2-CTD wild type as well as the mutants F743I and F743I/R747C were performed as described (Noell *et al.*, 2019). For the assays, His_6_-tagged *Hs* BicD2-CTD fragments (wild type and mutants) were purified as described by a single affinity chromatography step from 1L of cell culture, whereas Rab6^GTP^-GST and Nup358-min-GST where purified from 0.5L of cell culture. (Noell *et al.*, 2019). For GST-pull-down with Nup358-min, human Nup358 (residues 2147-2240) was purified by glutathione sepharose as described (Noell *et al.*, 2019) but not eluted. We then proceeded with the same protocol as described for the Rab6^GTP^ GST-pull-down (Noell *et al.*, 2019), however, GTP was omitted.

### Crystallization

Purified *Dm* BicD-CTD/F684I was set up for crystallization at 20°C in hanging drops. For the drop, 1 µl of the protein sample at a concentration of 8 mg/ml was mixed with 1 µl of reservoir buffer (4% PEG 3350, 0.4 M NaSCN, 5% glycerol). Crystals in space group *P*3_1_21 were obtained after 2-3 days in the dimensions 0.2 mm * 0.2 mm * 0.2 mm. Crystals were soaked in a cryo-buffer consisting of the reservoir solution with addition of 30% glycerol and 10 mM HEPES pH 7.5 and flash frozen in the liquid nitrogen.

### Structure determination

Data was collected from a single crystal at NE-CAT beam line 24ID-C at the Advanced Photon Source (APS), Argonne National Lab (ANL), which was equipped with a Pilatus 6M detector. X-ray intensities were processed and scaled using the RAPD software developed by F. Murphy, D. Neau, K. Perry and S. Banerjee, APS (https://rapd.nec.aps.anl.gov/login/login.html). The structure was determined by molecular replacement in the PHENIX suite (Adams *et al.*, 2010), with the structure of the wild-type as the search model, which was truncated N-terminal of residue 692 (Liu *et al.*, 2013). An initial model was obtained from automatic model building in the PHENIX suite and completed by manual model building in the program COOT (Adams *et al.*, 2010; Emsley *et al.*, 2010). The structure was refined through iterative cycles of manual model building and refinement (Adams *et al.*, 2010; Emsley *et al.*, 2010) to 2.35Å resolution in the PHENIX suite (Adams *et al.*, 2010), with an R_free_ of 25.99% and an R_work_ of 25.06% (Table 1). The stereochemical quality of the model was assessed with MolProbity (Chen *et al.*, 2010). The crystallographic statistics are summarized in Table 1.

### Structural analysis

Structures were compared by least-squares superimposition of the coordinates in COOT (Emsley *et al.*, 2010). Dimer interfaces and knobs-into-holes interactions were analyzed by the web servers PISA and SOCKET, respectively (Krissinel & Henrick, 2007; Walshaw & Woolfson, 2001). For identification of knobs-into-hole interactions, a cutoff of 7.5 Å was used (helix extension 1 residue). Figures were created in the PyMOL Molecular Graphics System, Version 2.0 (Schrödinger, LLC) and VMD (Humphrey *et al.*, 1996). The program APBS was used to analyze surface electrostatic potentials of proteins (Jurrus *et al.*, 2018). Default parameters were used; neutral charges were assigned to the N- and C-termini, waters were removed and selenomethionine residues were converted to methionine.

### Single molecule processivity assays with full-length *Dm* BicD

Dynein, dynactin and *Dm* BicD constructs were each diluted into 30 mM HEPES pH 7.4, 300 mM KOAc, 2 mM MgOAc, 1 mM EGTA, 20 mM DTT, clarified for 20 min at 400,000 x g and the concentration determined with the Bradford reagent (Bio-Rad). To form the dynein-dynactin-BicD (DDB) complex, dynein, dynactin and BicD (WT, CC1, or F684I) were incubated at a molar ratio of 1:1:2 (200 nM dynein, 200 nM dynactin and 400 nM BicD) on ice for 30 min in motility buffer (30 mM HEPES pH 7.4, 150 mM KOAc, 2 mM MgOAc, 1 mM EGTA, 2 mM MgATP, 20 mM DTT, 8 mg/ml BSA, 0.5 mg/ml kappa-casein, 0.5% pluronic F68, 10 mM paclitaxel and an oxygen scavenger system (5.8 mg/ml glucose, 0.045 mg/ml catalase, and 0.067 mg/ml glucose oxidase). To label the biotin-tag at the N-terminal region of the dynein heavy chain, 400 nM streptavidin-conjugated 655 quantum dots (Invitrogen) were added to the DDB complex and incubated on ice for 15 min. The DDB complex was diluted in motility buffer containing 50 mM KOAc to a final concentration of 0.5 nM dynein for observing motion on microtubules.

PEGylated slides were coated with 0.3 mg/ml rigor kinesin for attachment of rhodamine-labeled microtubules as described in (Sladewski *et al.*, 2018). Motility assays for all three DDB complexes (BicD^WT^, BicD^CC1^ and BicD^F684I^) were performed on three lanes of a single slide. Total Internal Reflection Fluorescence (TIRF) microscopy was used to capture images of Qdot labeled-dynein and microtubule tracks. Imaging was performed on a Nikon ECLIPSE Ti microscope equipped with through-objective type TIRF and run by the Nikon NIS Elements software. Images were captured at 200 ms temporal and 6 nm spatial resolution. Rhodamine-labeled microtubules and Qdot (655nm)-labeled dynein were excited with the 488 and 561 nm laser lines, respectively, and images simultaneously recorded at five frames/s using two Andor EMCCD cameras (Andor Technology USA, South Windsor, CT). Run-length was total travel distance, and speed was total travel distance divided by time. Binding frequency was normalized to number of events per time per µm microtubule. For statistical significance, an unpaired t-test with 95% confidence interval was performed for the speed data. For run length data, the Kolmogorov-Smirnov test with a 95% confidence interval was performed.

### Molecular dynamics simulations

MD simulations with implicit solvent were carried out using the CPU implementation of the PMEMD program in the AMBER16 package (Case *et al.*, 2016) as described (Noell *et al.*, 2019). The use of an implicit solvent model was justified by comprehensive comparisons of the results to those from explicit solvent simulations. The PMF or free energy associated with registry shift for the *Dm* BicD-CTD/F684I mutant was calculated using Eqn. (1) from a single 250 ns trajectory in which both the homotypic and asymmetric registries were sampled.

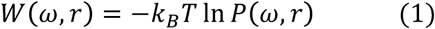

Here, *W* is the PMF as a function of a dihedral angle *ω*, more specifically, the C-C_α_-C_β_-C_γ_ dihedral angle of F691 of chain A, and a distance *r*, more specifically, the distance between the sidechain N atom of K678 of chain A and the C_δ_ atom of E673 of chain B (see Fig. 5). *k*_*B*_ is the Boltzmann constant, *T* is temperature, and *P* is a two-dimensional probability distribution function.

### CD spectroscopy

Purified *Dm* BicD-CTD/F684I (0.3 mg/ml) was dialyzed in the following buffer: 150 mM NaCl, 10 mM Tris pH 8 and 0.2 mM TCEP. CD spectra were recorded with a Jasco J-810 CD Spectrometer equipped with a thermoelectric control device, using a cuvette with a path length of 0.1cm. After the buffer baseline subtraction, CD signals were normalized to the protein concentration and converted to mean residue molar ellipticity [Θ]. CD spectra from 190 to 260nm were measured at 4°C or 90°C as described (Noell *et al.*, 2019). Thermal unfolding profiles of proteins were recorded by CD spectroscopy at 222 nm as described (Noell *et al.*, 2019).

## ACKNOWLEDGMENTS

This paper was supported by the following grants: National Institute of Health, National Institute of General Medical Sciences (NIH NIGMS) grant 1R15GM128119-01 awarded to S.R.S. and GM078097 to KMT. Additional funds came from the Chemistry Department at State University of New York at Binghamton and the Research Foundation of the State University of New York.

X-ray diffraction data was collected at Northeastern Collaborative Access Team (NE-CAT) beam line 24ID-C at the Advanced Photon Source (APS), Argonne National Lab (ANL), which are funded by the National Institute of General Medical Sciences from the National Institutes of Health (P30 GM124165) and by a NIH-ORIP HEI grant (S10 RR029205). APS is operated by ANL for the DOE under contract No. DE-AC02-06CH11357. We thank Jonathan P Schuermann from APS and Richard Vallee from Columbia University for helpful discussions. We also thank Carol Bookwalter and Elena Krementsova for cloning, protein expression and protein purification. This work used the Extreme Science and Engineering Discovery Environment (XSEDE)(Towns *et al.*, 2014), which is supported by NSF grant ACI-1548562. All figures related to the computational studies were made using the Visual Molecular Dynamics (VMD) program (Humphrey *et al.*, 1996). Furthermore, we thank S. Bane & B. Callahan (SUNY Binghamton), for access to equipment. Coordinates and structure factors were deposited to the protein data bank (https://www.wwpdb.org, under PDB ID 6TZW). The content is solely the responsibility of the authors and does not necessarily represent the official views of the National Institutes of Health.

