## Supplementary Information for "Coiled-coil registry shifts in the F684I mutant of Bicaudal result in cargo-independent activation of dynein motility"

### SUPPORTING INFORMATION

**Table of Contents** **Page**

**Supporting Methods** S2

**Table S1.** Dimer interface of crystal structure of *Dm* BicD-CTD/F684I. S3

**Figure S1.** Assignment of heptad repeats for BicD-CTD F684I. S4

**Figure S2.** The composition of the crystallization buffers does not affect the α-helical content of *Dm* BicD-CTD WT or the F684I mutant. S5

**Figure S3.** MD simulations suggest that a conformation of human BicD2-CTD/F743I mutant with homotypic registry is stable. S6

**Supporting Movie Legends** (Movies S1, S2 and S3). S7

**Description of Other Supporting Files** (Files S1 – S5). S7

**References** S8

### SUPPORTING METHODS

#### CD spectroscopy: thermal unfolding profiles

All proteins revealed a sigmoidal denaturation profile, characteristic of a two-state transition from helix to random coil, with a single melting temperature ($T_M$), therefore, a two-state denaturation model (α-helix to random coil) was used to analyze the data. The apparent melting temperature ($T_M$) was determined from the peak of smoothened differential melting curves d[Θ222](T)/dT using Origins Pro software and averaged from three experiments. Representative melting curves are shown. The fraction of folded peptide ($F_{folded}$) was calculated from the equation $F_{folded} = ([\Theta]-[\Theta]_{unfolded})/([\Theta]_{folded}-[\Theta]_{unfolded})$, where [Θ] is the observed molar ellipticity at any particular temperature and $[\Theta]_{unfolded}$ and $[\Theta]_{folded}$ are the molar ellipticities of the denatured (unfolded) and native (folded).

### SUPPORTING TABLES

**Table S1 Dimer interface of crystal structure of *Dm* BicD-CTD/F684I. For the corresponding Table for *Hs* BicD2-CTD see (Noell *et al.*, 2019).**

| ***Dm* BicD-CTD F684I** | | | ***Dm* BicD-CTD** (PDB ID 4BL6) | | |
|---|---|---|---|---|---|
| Chain A | Chain B | Dist(Å) | Chain A | Chain D | Dist (Å) |
| **Hydrogen bonds** | | | **Hydrogen bonds** | | |
| TYR 698 OH | CYS 695 SG | 3.05 | TYR 698 OH | CYS 695 SG | 3.2 |
| GLN 701 OE1 | TYR 698 OH | 2.70 | GLN 701 OE1 | TYR 698 OH | 3.0 |
| LYS 716 NZ | GLU 715 OE1 | 3.42 | GLU 715 OE1 | LYS 716 NZ | 2.5 |
| ASN 720 ND2 | GLU 715 OE2 | 3.70 | ARG 688 NH1 | ASP 680 OD1 | 2.8 |
| | | | ARG 688 NH2 | THR 683 OG1 | 2.8 |
| **Salt bridges** | | | **Salt bridges** | | |
| GLU 715 OE1 | LYS 716 NZ | 3.37 | GLU 715 OE1 | LYS 716 NZ | 2.5 |
| GLU 715 OE2 | LYS 716 NZ | 3.29 | GLU 715 OE2 | LYS 716 NZ | 3.9 |
| LYS 716 NZ | GLU 715 OE1 | 3.42 | ARG 688 NH1 | ASP 680 OD1 | 2.8 |
| LYS 716 NZ | GLU 715 OE2 | 3.90 | ARG 688 NH2 | ASP 680 OD1 | 3.1 |
| | | | LYS 716 NZ | GLU 715 OE2 | 2.9 |
| **Interface area** | | | **Interface area** | | |
| 1427 Å$^2$ | | | 1764 Å$^2$ | | |
| **Interface residues** | | | **Interface residues** | | |
| 684, 685, 687, 688, 691, 694, 695, 698, 701, 702, 705, 706, 708, 709, 712, 713, 715, 716, 719, 720, 722, 723, 725, 726, 727, 729, 730, 733, 734, 736, 737, 740 | | | 659, 662, 663, 664, 666, 667, 669, 670, 671, 673, 674, 677, 678, 680, 681, 683, 684, 687, 688, 690, 691, 692, 694, 695, 698, 699, 701, 702, 705, 706, 708, 709, 712, 713, 715, 716, 719, 720, 722, 723, 726, 727, 729, 730, 733, 734, 736, 737, 740 | | |

Standard nomenclature from the structure coordinates is used for Chain ID, residue and atom names. Dist: Distance. Analysis was performed with the PISA server (Krissinel & Henrick, 2007).

**SUPPORTING FIGURES**

### (A) Assignment of heptad repeats for *Dm* BicD-CTD F684I

```
Chain A
MSKLRNELRLLKEDAATISSLRAMFAARCEEYVTQVDDLNRQLEAAEEEKKTLNQLLRLAVQQKLALTQRLEE
Register---------------------------abcdefgabcdefgabcdefgabcdefgabcdefga--
Knobs------------------------------Y--Y---Y--Y---Y--Y---Y--Y---Y--Y------

Chain B
--------------------LRAMFAARCEEYVTQVDDLNRQLEAAEEEKKTLNQLLRLAVQQKLALTQRLEE
Register---------------------------abcdefgabcdefgabcdefgabcdefgabcdefga--
Knobs---------------X---X-------------X---X--X---X--X---X--X---X--X------
```

### (B) Assignment of heptad repeats for *Dm* BicD-CTD (PDB ID 4BL6)

```
Chain A
-----VSDTMSKLRNELRLLKEDAATFSSLRAMFAARCEEYVTQVDDLNRQLEAAEEEKKTLNQLLRLAVQQKLALTQRLEEM
                abcdefgabcdefgabcdefgabcdefgabcdefgabcdefgabcdefgabcdefgabcdefga
------------Y---Y--Y----------Y-------------Y--Y---Y--Y---Y--Y---Y--Y------Y-------

Chain D
NEKIIVSDTMSKLRNELRLLKEDAATFSSLRAMFAARCEEYVTQVDDLNRQLEAAEEEKKTLNQLLRLAVQQKLALTQRLEEM
     abcdefgabcdefgabcdefgabcdefgabcdefgabcdefgdefgabcdefgabcdefgabcdefgabcdefga
-----X--X---X--X----------X------X-------------X---X--X---X--X---X---------X-------
```

**Figure S1 Assignment of heptad repeats for BicD-CTD F684I.** (A, B) The protein sequences from the structures are shown and the Chain IDs from the coordinates are indicated. Y and X denote knob residues that form "knobs-into-holes" interactions with the other chain of the dimer. Knobs-into-holes interactions were identified in the structures with the program SOCKET (Walshaw & Woolfson, 2001). A cutoff of 7.5Å and a helix extension of 1 residue were used. Knob residues are denoted by 'Y' and 'X' and are located at 'a' and 'd' positions of the heptad repeat. The unedited output from SOCKET is shown. In Figure 1, all 'a' position residues are shown in sphere representation for which at least one knob was confirmed by SOCKET in a layer. To clarify, if the second 'a' position residue of the layer was not identified as knob by SOCKET, it is still shown in sphere representation, consistent with references (Liu *et al.*, 2013; Terawaki *et al.*, 2015). See also (Noell *et al.*, 2019). For the corresponding figure of *Hs* BicD2-CTD and sequence alignments of *Dm* BicD, *Ms* BicD1 and *Hs* BicD2 see (Noell *et al.*, 2019).

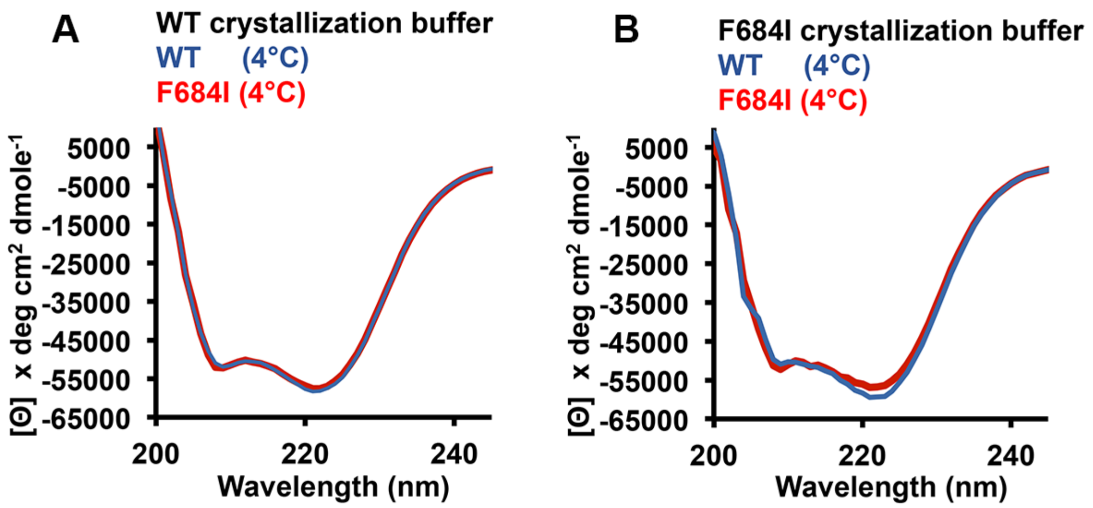


**Figure S2. The composition of the crystallization buffers does not affect the α-helical content of *Dm* BicD-CTD WT or the F684I mutant.** (A, B) In the structure of the *Dm* BicD-CTD/F684I mutant, a ~20 residue region is disordered, whereas this region is α-helical in the wild-type structure. In order to exclude the possibility that differences in the chemical composition of the WT and F684I mutant crystallization buffers were responsible for the observed structural differences, we recorded CD wavelength scans of *Dm* BicD-CTD WT (blue) and *Dm* BicD-CTD/F684I mutant (red) at 4°C in (modified) crystallization buffers. The mean residue molar ellipticity [Θ] versus the wavelength is shown. The buffers were modified for CD spectroscopy in order to reduce the background and to prevent crystallization of the proteins. Note that the α-helical content of the *Dm* BicD-CTD WT and F684I proteins is very similar in both crystallization buffers (see also Fig. 4B). Based on these data, it is unlikely that the disorder in the ~20 residue region that is observed in the structure of the *Dm* BicD-CTD/F784I mutant is caused by differences in the chemical composition of the crystallization buffers. Therefore, we conclude that the observed disorder is likely caused by structural flexibility rather than unfolding. (A) The following buffer was used for crystallization of the *D*m BicD-CTD WT: 0.1 M Tris pH 8.5, 5% PEG4000, 30 mM $CaCl_2$, 30 mM $MgCl_2$, 10% glycerol, 10 – 20 mM HEPES pH 7.4, 75 - 150 mM NaCl, and 2.5 - 5 mM β-mercapthoethanol (BME). CD spectroscopy was performed in a modified version of this buffer: 20 mM Tris pH 8.5, 2.5% PEG4000, 15 mM $CaCl_2$, 15 mM $MgCl_2$, 75 mM NaCl, and 0.2 mM TCEP. (B) The following buffer was used for crystallization of the *Dm* BicD-CTD/F743I mutant: 4% PEG3350, 0.4 M NaSCN, 5 % glycerol, 10-20mM HEPES pH 7.5, 75-150 mM NaCl, and 0.2-0.5 mM TCEP. CD spectroscopy was performed in a modified version of this buffer: 10 mM Tris pH 7.5, 3 mM NaSCN, 150mM NaCl, and 0.2mM TCEP.

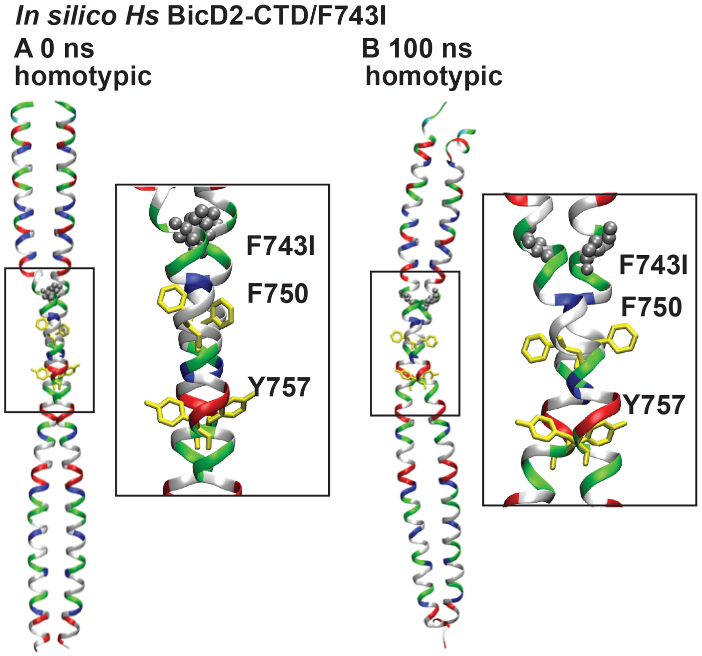


**Figure S3 MD simulations suggest that the human BicD2-CTD/F743I mutant can assume a conformation with a homotypic coiled-coil registry.** For these simulations, the structure of *Hs* BicD2-CTD was chosen as a starting point, since it has a homotypic coiled-coil registry (Noell *et al.*, 2019), and the homologous F743I mutation was introduced. **(**A) Cartoon representation of the crystal structure of human BicD2-CTD (Noell *et al.*, 2019), with homotypic registry, colored by residue type (blue: positively charged, red: negatively charged, green: polar, white: non-polar). F743 was mutated to isoleucine (silver spheres). F750 and F757 are shown in yellow stick representation. A close-up of the boxed area is shown on the right. See also File S4. (B) Equilibrated structure of the F743I mutant of human BicD2-CTD at the end of a 100 ns MD simulation. Note that in these MD simulations, the homotypic registry of the *Hs* BicD2-CTD/F743I was preserved, suggesting that the mutant can assume a conformation with a homotypic registry. See also File S5. In comparison, in our recent MD simulations of *Hs* BicD2-CTD/F743I with asymmetric coiled-coil registry, the F743I mutation induced a coiled-coil registry shift from an asymmetric to a fully heterotypic registry (Noell *et al.*, 2019), suggesting that the human homolog of the Bicaudal mutant can also sample multiple conformations with distinct coiled-coil registries.

### SUPPORTING MOVIE LEGENDS

**Movie S1** Diffusive motion of dynein-dynactin-BicD WT ($DDB^{WT}$) on rhodamine-labeled microtubules. The dynein is labeled with a Qdot. Movies are 2x real speed. Horizontal distance, 20 µm.

**Movie S2** Processive motion of dynein-dynactin-BicD-CC1 ($DDB^{CC1}$) on rhodamine-labeled microtubules. The dynein is labeled with a Qdot. Movies are 2x real speed. Horizontal distance, 20 µm.

**Movie S3** Processive motion of dynein-dynactin-BicD/F684I ($DDB^{F684I}$) on rhodamine-labeled microtubules. The dynein is labeled with a Qdot. Movies are 2x real speed. Horizontal distance, 20 µm.

### DESCRIPTION OF OTHER SUPPORTING FILES

**File S1** Equilibrated coordinates of *Dm* BicD-CTD/F684I, with homotypic coiled-coil registry. F684 was mutated to isoleucine (see Fig. 5A).

**File S2** Coordinates of the F684I mutant of *Dm* BicD-CTD after ~53 ns of an MD simulation (see Fig. 5B).

**File S3** Coordinates of the F684I mutant of *Dm* BicD-CTD after ~120 ns of the same MD simulation. (see Fig. 5C).

**File S4** Initial coordinates for MD simulations of *Hs* BicD2-CTD/F743I (see Fig. S3A).

**File S5** Final equilibrated coordinates for MD simulations of *Hs* BicD2-CTD/F743I (see Fig. S3B).
